# Centrosome maturation requires phosphorylation-mediated sequential domain interactions of SPD-5

**DOI:** 10.1101/2021.05.20.444955

**Authors:** Momoe Nakajo, Hikaru Kano, Kenji Tsuyama, Nami Haruta, Asako Sugimoto

**Affiliations:** Laboratory of Developmental Dynamics, Graduate School of Life Sciences, Tohoku University, 2-1-1 Katahira, Aoba-ku, Sendai, 980-8577, Japan

## Abstract

Centrosomes consist of two centrioles and surrounding pericentriolar material (PCM). PCM expands during mitosis in a process called centrosome maturation, in which PCM scaffold proteins play pivotal roles to recruit other centrosomal proteins. In *C. elegans*, the scaffold protein SPD-5 forms PCM scaffold in a PLK-1 phosphorylation-dependent manner. However, how phosphorylation of SPD-5 promotes PCM scaffold assembly is unclear. Here, we identified three functional domains of SPD-5 through *in vivo* domain analyses, and propose that sequential domain interactions of SPD-5 are required for mitotic PCM scaffold assembly. Firstly, SPD-5 is targeted to centrioles through direct interaction between its centriole localization (CL) domain and a centriolar protein PCMD-1. Then, intra- and inter-molecular interaction between SPD-5 phospho-regulated multimerization (PReM) domain and the PReM association (PA) domain is triggered by phosphorylation by PLK-1, which leads to PCM scaffold expansion. Our findings suggest that the sequential domain interactions of scaffold proteins mediated by Polo/PLK-1 phosphorylation is an evolutionarily conserved mechanism of PCM scaffold assembly.

## Introduction

Centrosomes, the major microtubule organizing center (MTOC) in animal cells, consist of two centrioles and surrounding amorphous mass of proteins called pericentriolar material (PCM). The size of PCM dramatically changes in a cell-cycle dependent manner. During interphase, a small amount of PCM is assembled adjacent to the centrioles. When cells enter mitosis, centrosomal proteins including γ-tubulin complexes are recruited to the PCM to expand its size and the microtubule nucleation activity is increased. This process is called centrosome maturation (Palazzo et al., 2000).

During centrosome maturation, PCM scaffold proteins play pivotal roles to recruit other centrosomal proteins around centrioles. One of the well characterized PCM scaffold proteins is Centrosomin (Cnn) in *Drosophila melanogaster* and its ortholog CDK5RAP2 in humans (Fong et al., 2008; Megraw et al., 1999). In *C. elegans*, SPD-5 is thought to be a functional homolog of Cnn/CDK5RAP2, while sequence homology is undetectable (Hamill et al., 2002). CDK5RAP2/Cnn/SPD-5 contain multiple predicted coiled-coil regions, and scaffold assembly of these proteins are facilitated by the interaction with the conserved coiled-coil proteins Cep192/SPD-2 and phosphorylation by Polo/PLK-1 (Conduit et al., 2014a; Decker et al., 2011; Giansanti et al., 2008; Gomez-Ferreria et al., 2007; Haren et al., 2009; Kemp et al., 2004; Pelletier et al., 2004). These similarities suggest that a universal mechanism of PCM scaffold assembly may exist.

Through the studies of Cnn, its distinct domains and their roles in scaffold assembly have been revealed. Cnn have two motifs that are conserved in CDK5RAP2; the N-terminal CM1 (Centrosomin motif 1) domain that recruits γ-tubulin complexes to the centrosomes (Fong et al., 2008; Zhang and Megraw, 2007), and the C-terminal CM2 domain that is implicated in their centrosomal targeting (Barr et al., 2010; Feng et al., 2017). During early mitosis, Cnn is recruited to the centrioles mainly by Spd-2, then phosphorylated by Polo at the central region called the PReM (Phospho-regulated multimerization) domain (Alvarez-Rodrigo et al., 2019; Conduit et al., 2014a; Conduit et al., 2014b). The phosphorylation of the PReM domain enhances the interaction between this domain and the CM2 domain. This phosphorylation-dependent conformational change of Cnn leads to the assembly of the PCM scaffold (Feng et al., 2017).

In *C. elegans*, PCM scaffold protein SPD-5 is recruited to centrioles by centriolar/PCM protein SPD-2, and they depend on each other for localization to the expanded PCM (Hamill et al., 2002; Kemp et al., 2004; Pelletier et al., 2004). SPD-2 also function as a PLK-1 recruiter, and PLK-1 phosphorylation activity is required for the centrosome maturation (Decker et al., 2011; Erpf et al., 2019). The mutation of PLK-1 phosphorylation sites of SPD-5 central region results in the failure of the scaffold assembly *in vivo* and *in vitro* (Woodruff et al., 2015; Wueseke et al., 2016). *In vitro* reconstitution studies further demonstrated that the purified SPD-5 assembles into spherical condensates which can recruit other downstream PCM proteins. This SPD-5 scaffold assembly is robustly promoted by addition of SPD-2 and PLK-1 (Woodruff et al., 2017; Woodruff et al., 2015). Recently, the centriolar/PCM protein PCMD-1 was identified as a novel factor involved in PCM organization. Genetic analyses indicated that PCMD-1 required for recruitment of SPD-5 around centrioles and the integrity of PCM scaffold (Erpf et al., 2019). Although the involvement of SPD-2, PCMD-1 and PLK-1 is implicated, how SPD-5 interacts with each of these proteins and is regulated by phosphorylation in the PCM scaffold assembly is poorly understood.

In this study, we conducted *in vivo* domain analysis of SPD-5, and identified three functional domains. We propose a model for the SPD-5 scaffold assembly during centrosome maturation through sequential interactions of these domains mediated by PLK-1-dependent phosphorylation.

## Results

### Regions in the C-terminal half of SPD-5 are involved in PCM localization

To understand the mechanism(s) by which SPD-5 is localized to the centrosomes and participates in PCM scaffold formation, we conducted a domain analysis of SPD-5 *in vivo*. Several truncated versions of SPD-5 proteins tagged with green fluorescent protein (GFP) at their N terminus were expressed in early *C. elegans* embryos from a transgene inserted into the specific locus on Chromosome II that is distinct from the *spd-5* locus. The localization of transgenic proteins and embryonic phenotypes were investigated in the presence and the absence of endogenous SPD-5. The expression of each SPD-5 fragment was confirmed by western blotting of worm extracts (Fig. S1).

In both the presence and absence of endogenous SPD-5, the full-length SPD-5 (GFP::SPD-5(FL); amino acids [aa] 1–1198) localized to the centrosomes throughout the cell cycle in all observed early embryos, as reported (Woodruff et al., 2015) (n = 15, and 21, respectively) (Fig. 1A). As GFP::SPD-5(FL) fully rescued the embryonic lethality of *spd-5(RNAi)* (n = 21), we concluded that this fusion protein is functional *in vivo*. However, we noted that, in the absence of the endogenous SPD-5, the centrosomal signal was weaker and its shape was slightly distorted, which might be caused by the lower amount of GFP::SPD-5(FL) relative to that of endogenous SPD-5 (Fig. S1) and/or the effect of the GFP tag.

**Figure 1 :**
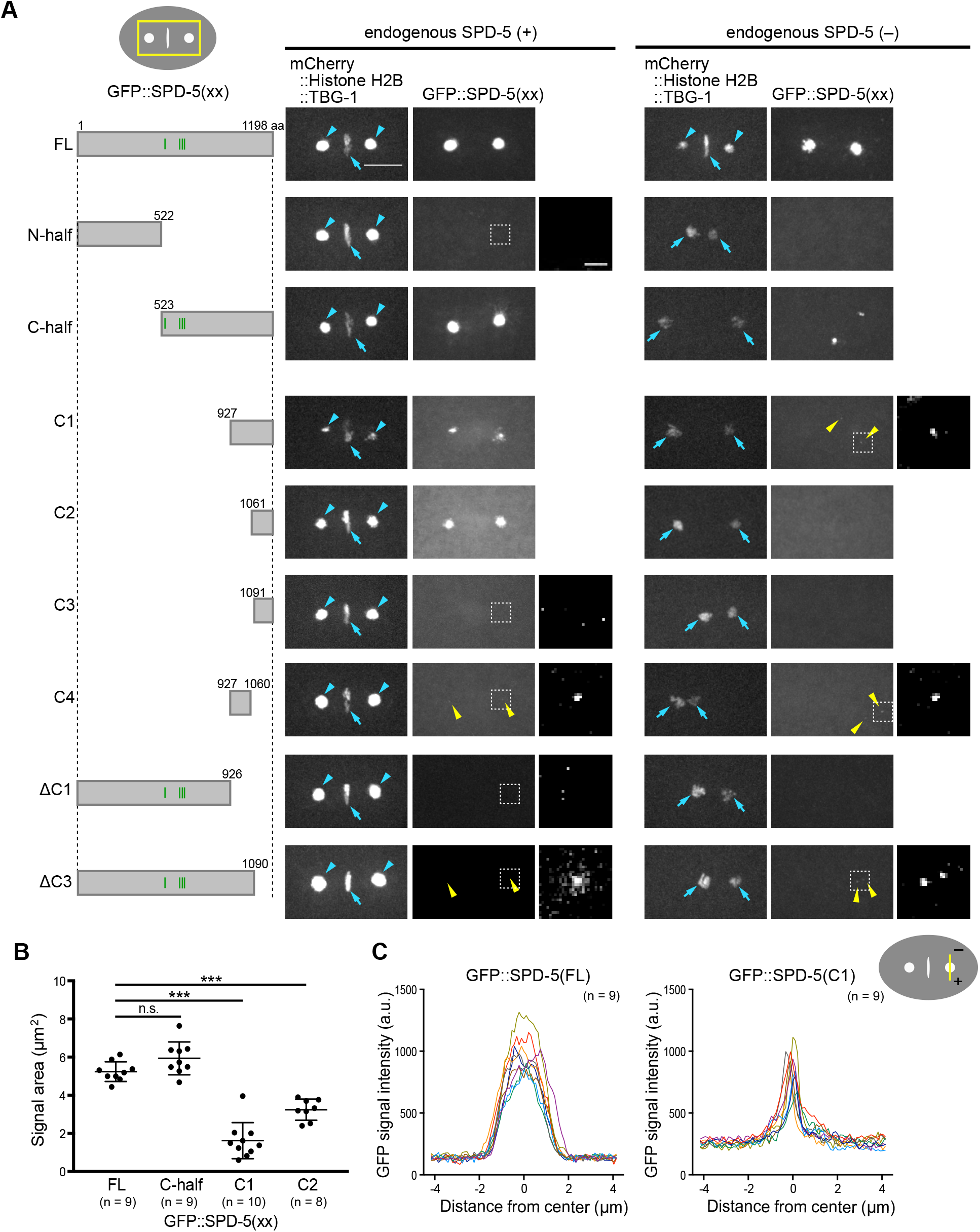
*In vivo* domain analysis of SPD-5. (A) Images of *C. elegans* one-cell embryos that express full-length or truncated GFP::SPD-5 proteins (GFP::SPD-5(xx)), mCherry::Histone H2B and mCherry::TBG-1/γ-tubulin. The gray boxes in the left column indicate the fragments used in the assay. The four green vertical lines indicate the serine residues that can be phosphorylated by PLK-1 (S530, S627, S653 and S658) (Woodruff et al., 2015). The areas around the centrosomes and chromosomes, illustrated as a yellow rectangle in the upper-left diagram, are shown in each image. The images were taken at 160 sec after nuclear envelope breakdown, which corresponds to metaphase in the control embryos. Scale bar, 10 μm. Blue arrows indicate mCherry::Histone H2B (a chromosome marker), and blue arrowheads indicate mCherry::TBG-1 (a centrosome marker). Yellow arrowheads indicate the faint and small GFP signals on and around the centrioles. The magnified and enhanced images of the areas in the dotted white squares are shown in the right column. Scale bar, 1 μm. (B) Area of centrosomal GFP signals for each GFP::SPD-5(xx) in the presence of endogenous SPD-5 at metaphase. The mean ± SEM is indicated. ***p < 0.001; n.s., not significant (p ≥ 0.05), Tukey’s multiple comparisons test. (C) Signal distribution of each GFP::SPD-5(xx) at centrosomes in the presence of endogenous SPD-5. Signals were measured along the short axis of the embryo, illustrated with a yellow line in the schematic (upper right). a.u., arbitrary units.

We first divided the SPD-5 protein into two regions, the N-terminal half (N-half, aa 1–522) and the C-terminal half (C-half, aa 523–1198) (Fig. 1 A). In all observed embryos, GFP::SPD-5(N-half) signal was barely detected at centrosomes in either the presence (n = 10) or absence (n = 5) of endogenous SPD-5, although in the latter case some condensates were detected in the cytosol at prometaphase to anaphase (data not shown).

In contrast, GFP::SPD-5(C-half) localized to the centrosome in the presence of endogenous SPD-5 (100%, n = 11) (Fig. 1 A). In the absence of endogenous SPD-5, the centrosomal signal of GFP::SPD-5(C-half) became weaker and smaller, indicative of its centriolar localization (100%, n = 15). These data indicate that GFP::SPD-5(C-half) can interact with endogenous SPD-5 to localize to the centrosomes, but it is deficient for the expansion of the PCM. These results suggest that the C-terminal half region of SPD-5 is involved in its targeting to the centrioles and interacts with endogenous SPD-5.

### SPD-5(C1) has a dominant-negative effect on PCM scaffold formation by endogenous SPD-5

We further dissected the SPD-5 C-terminal half region for its ability to interact with SPD-5 itself and centriolar targeting. When the C-terminal 272 aa (C1 region, aa 927–1198) were deleted (GFP::SPD-5(ΔC1)), localization to the PCM and the centrioles was abolished in either the presence (100%, n = 4) or absence (100%, n = 5) of endogenous SPD-5 (Fig. 1 A), suggesting that the C1 region is necessary for both self-interaction and centriolar localization.

In turn, the C1 region (GFP::SPD-5(C1)) localized to the centrosome in the presence of endogenous SPD-5 (100%, n = 9; Fig. 1 A), but the signal area was significantly smaller (31% of that of GFP::SPD-5(FL)) and was fragmented (Fig. 1 B and C). Immunostaining using an antibody against SPD-5 (anti–SPD-5) that recognizes both endogenous SPD-5 and GFP::SPD-5(C1) revealed that endogenous SPD-5 co-localized with the disordered and fragmented signal of GFP::SPD-5(C1) (Fig. 2 A). Notably, this strain showed 46% embryonic lethality (Fig. 2 B). These results suggest that the C1 fragment disrupts ability of endogenous SPD-5 to assemble the PCM scaffold in a dominant-negative manner.

**Figure 2 :**
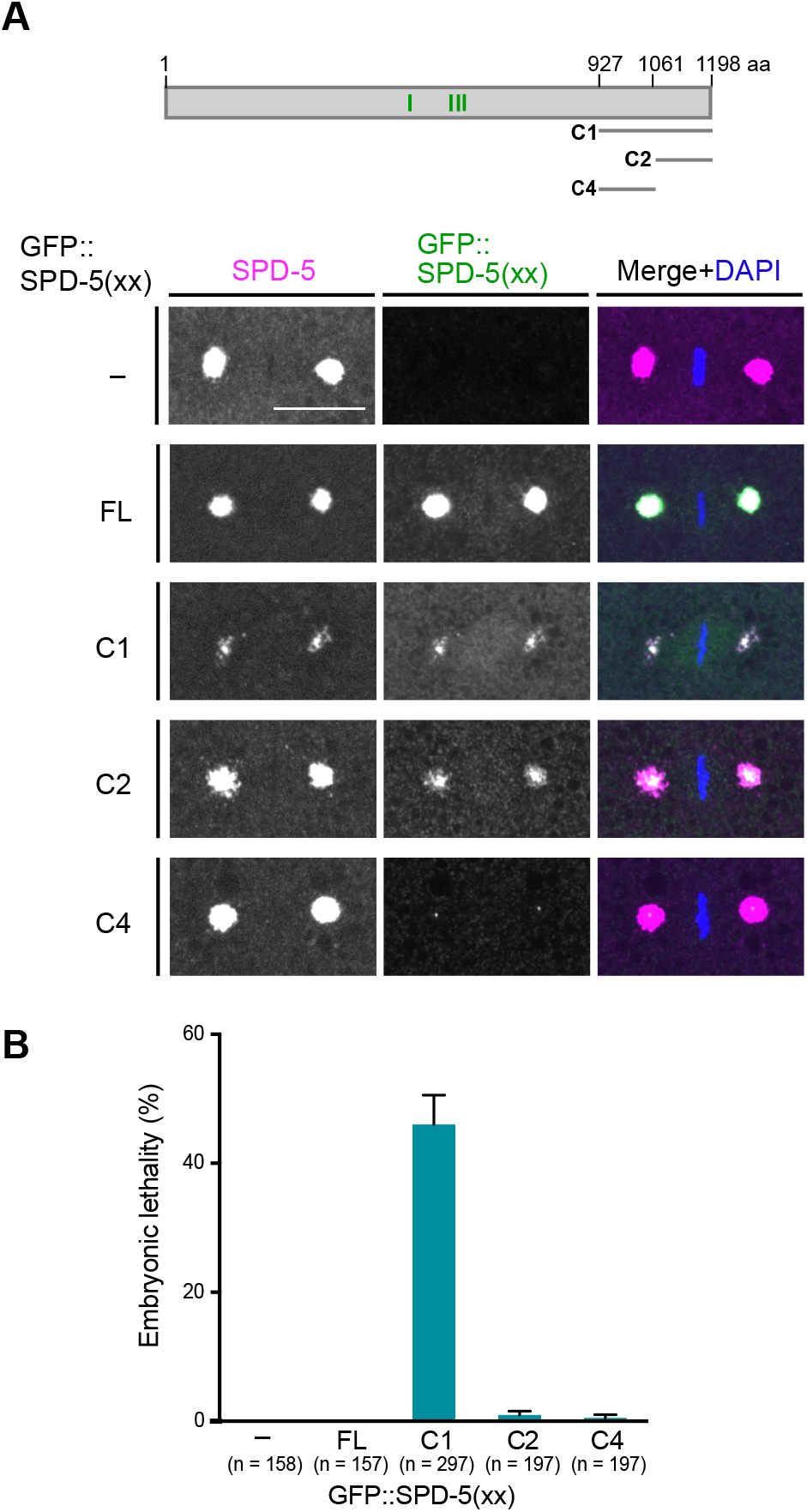
GFP::SPD-5(C1) has a dominant-negative effect on PCM scaffold formation by endogenous SPD-5. (A) Immunostaining of wild-type embryos and embryos expressing GFP::SPD-5(xx) using anti–SPD-5 and anti-GFP. The top diagram indicates the fragments used. The four green vertical lines indicate serine residues that can be phosphorylated by PLK-1. The SPD-5 antibody stains both endogenous SPD-5 and each GFP::SPD-5(xx). Scale bar indicates 10 μm. (B) Embryonic lethality of each GFP::SPD-5(xx) expressing strain. Error bars indicate +SEM.

In the absence of endogenous SPD-5, the GFP::SPD-5(C1) signal became even weaker and smaller, implying that this region of SPD-5 was localized only around the centrioles. These results suggest that the C1 region is involved in centriolar targeting and in intermolecular interactions with other SPD-5 molecules.

### The C-terminal region of SPD-5 contains two functional domains

The C-terminal half of the C1 region contains a region that is highly conserved among the *Caenorhabditis* genus, which we refer to as the C2 region (aa 1061–1198). In the presence of endogenous SPD-5, GFP::SPD-5(C2) was localized to the centrosome (100%, n = 8), although the signal area was 38% smaller than that of SPD-5(FL) (Fig. 1 A and B). A further deletion of 30 aa (C3, aa 1091–1198) abolished this centrosomal localization (100%, n = 5; Fig. 1 A). Unlike GFP::SPD-5(C1), GFP::SPD-5(C2) did not show a dominant-negative effect, and the area of endogenous SPD-5 was unaffected (Fig. 2 A and B). This centrosomal localization of GFP::SPD-5(C2) was lost in the absence of SPD-5 (92%, n = 12; Fig. 1 A). Thus, the C2 region interacts with endogenous SPD-5 in the central region of the PCM, but it lacks the ability to independently localize around the centrioles.

The fragment lacking the majority of the C2 region (GFP::SPD-5(ΔC3)) in the presence of endogenous SPD-5 localized around the centrioles but failed to expand to the PCM (100%, n = 4; Fig. 1 A). This suggests that the C2 region is dispensable for centriolar localization but is necessary for incorporation into the PCM.

The remaining N-terminal half of the C1 region is referred to as the C4 region (aa 927–1060). In both the presence and absence of endogenous SPD-5, GFP::SPD-5(C4) localization was represented by very small signals, plausibly around the centrioles (92%, n = 13; 100%, n = 12, respectively) (Fig. 1 A), suggesting that the C4 region by itself can localize to the centrioles, independently of the interaction with endogenous SPD-5. GFP::SPD-5(C4) did not show a dominant-negative effect on PCM scaffold assembly of endogenous SPD-5 (Fig. 2 A and B).

Taken together, these results identified two functional regions in the C-terminal half of SPD-5: C2 (aa 1061–1198), which is involved in the intermolecular interaction with endogenous SPD-5, and C4 (aa 927–1198), which is necessary and sufficient for centriolar targeting.

### The C2 region of SPD-5 interacts with the phosphorylated SPD-5 central region

Next, we examined which region of SPD-5 interacts with the C2 region by yeast two-hybrid assays. The C2 region (and the C1 region, which contains C2) interacted with the middle region of SPD-5 (aa 272–732) (Fig. 3 A). The shorter fragments SPD-5(272–680), SPD-5(400–680) and SPD-5(523–732) did not interact with the C2 region (Fig. 3 A), implying that a relatively long (∼460 aa) central region is required for the interaction with the C2 region.

**Figure 3 :**
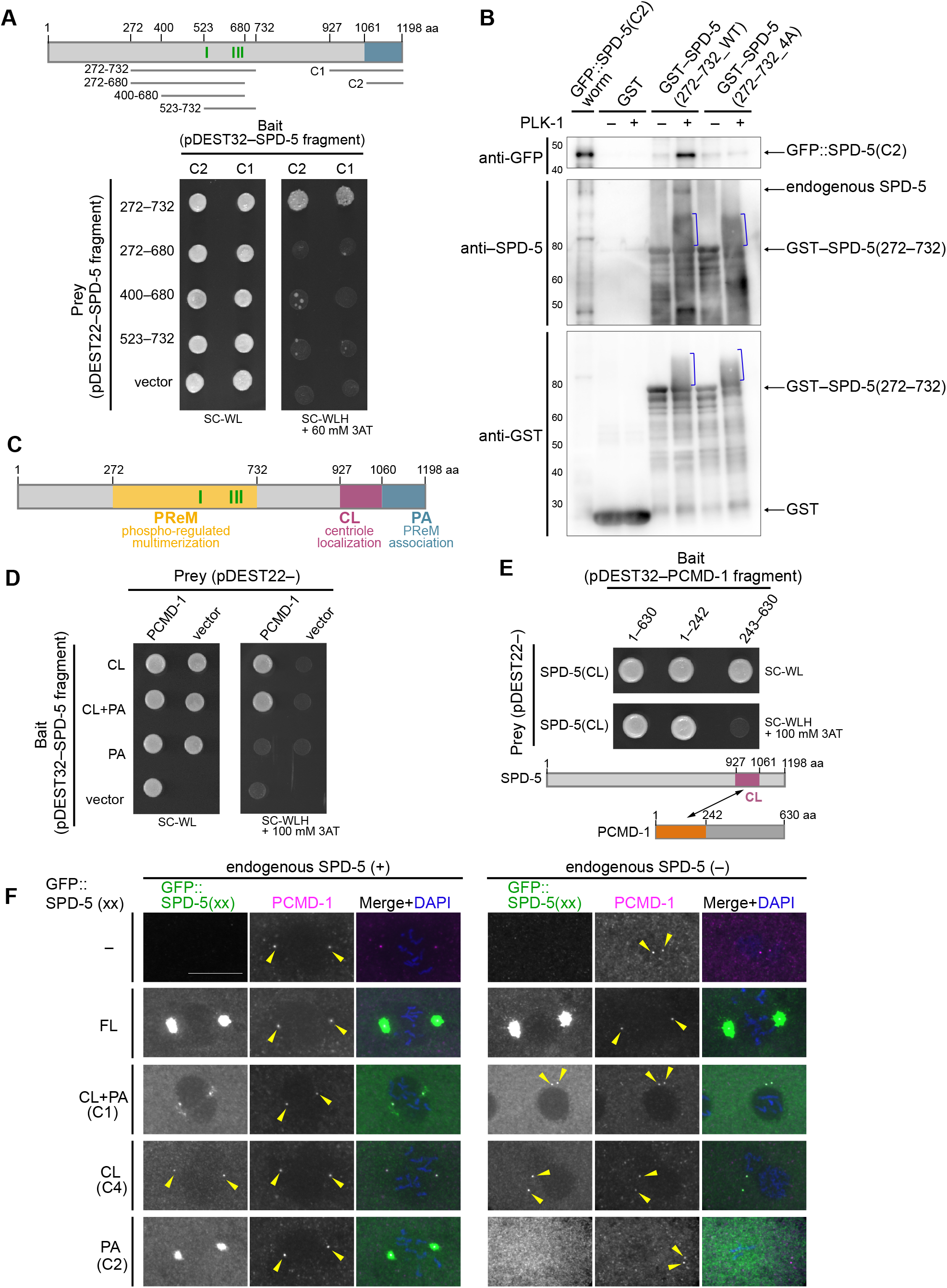
Identification of three functional domains of SPD-5 and their interactions. (A) Yeast two-hybrid analysis of interactions between SPD-5 fragments. The upper diagram shows the region corresponding to each SPD-5 fragment. The four green vertical lines indicate serine residues that can be phosphorylated by PLK-1. The growth of each yeast strain on SC/–Trp/–Leu plates (SC-WL, control) and SC/–Trp/–Leu /–His plates (SC-WLH) containing 60 mM 3 amino-1,2,4 triazol (3AT) is shown. (B) Pull-down assay for the phosphorylation-dependent interaction between SPD-5(C2) and SPD-5(272–732). Western blotting using anti-GFP, anti SPD-5 and anti-GST is shown. Preincubation of each GST-tagged protein with PLK-1 is indicated by +/−. Blue brackets correspond to phosphorylated GST SPD-5(272–732_WT) and GST SPD-5(272–732_4A) by PLK-1. Since SPD-5(272–732) has been shown to contain other phosphorylation sites than the mutated sites(Woodruff et al., 2015), SPD-5(272–732_4A) would still be phosphorylated by PLK-1. GFP::SPD-5(C2), 49.5 kDa; endogenous SPD-5, 135.1 kDa; GST SPD-5(272–732_WT), 78.2 kDa; GST SPD-5(272–732_4A), 78.1 kDa; GST, 26.8 kDa. (C) The three functional domains of SPD-5 identified in this study. (D and E) Yeast two-hybrid analysis of interactions between SPD-5 and PCMD-1. Growth on SC/–Trp/–Leu plates (SC-WL, control) and SC/–Trp/–Leu /–His plates (SC-WLH) containing 100 mM 3AT is shown. (D) Assessment of PCMD-1 interactions with SPD-5(CL), SPD-5(CL+PA) and SPD-5(PA). (E) Analysis of PCMD-1 interactions with SPD-5(CL). (F) Colocalization of SPD-5(xx) and PCMD-1 in one-cell embryos. Embryos were immunostained using anti-GFP and anti–PCMD-1. Images of one-cell embryos at the stage from pronuclear migration to prometaphase are shown. Yellow arrowheads indicate the small PCMD-1 or GFP signals on and around the centrioles. Scale bar indicates 10 μm.

As SPD-5(272–732) contains the PLK-1 phosphorylation sites required for expansion of the SPD-5 scaffold (Woodruff et al., 2015; Wueseke et al., 2016), we next examined by pull-down assay whether their phosphorylation by PLK-1 affects the interaction with SPD-5(C2). Purified glutathione *S*-transferase (GST)-tagged SPD-5(272–732) proteins with or without PLK-1 pretreatment were incubated with *C. elegans* egg lysates from the strain that expresses GFP::SPD-5(C2). The interaction of GST-tagged SPD-5(272–732_WT) with GFP::SPD-5(C2) and endogenous SPD-5 was detectable after pretreatment with PLK-1. In contrast, when the key PLK-1 phosphorylation sites (S530, S627, S653 and S658) were substituted with alanine, i.e., SPD-5(272–732_4A), its interaction with GFP::SPD-5(C2) and endogenous SPD-5 was barely detectable regardless of PLK-1 pretreatment (Fig. 3 B). These results imply that the PLK-1 phosphorylation within SPD-5(272–732) promotes the interaction with SPD-5(C2) and with the C2 region of endogenous SPD-5.

These *in vivo* and *in vitro* domain analyses revealed three functional regions of SPD-5 (Fig. 3 C). The C4 region was necessary and sufficient for centriolar targeting and hereafter is called the CL (centriole localization) domain. The SPD-5(272–732) region, which contains the key PLK-1 phosphorylation sites and was essential for SPD-5 scaffold expansion, is called the PReM (phospho-regulated multimerization) domain, which is named after the Centrosomin-PReM domain in *Drosophila* (Conduit et al., 2014a). The C2 region, which was necessary and sufficient for the phosphorylation-dependent interaction with SPD-5(PReM) is referred to as the PA (PReM association) domain (Fig. 3 C).

### SPD-5(CL) physically interacts with the centriolar protein PCMD-1

To identify proteins involved in the recruitment of the SPD-5(CL) domain to the centrioles, we conducted a yeast two-hybrid screen. Using SPD-5(CL+PA) as the bait, the centriolar protein PCMD-1 was identified; PCMD-1 also interacted with SPD-5(CL), but not with SPD-5(PA) (Fig. 3 D). SPD-5(CL) interacted with the N-terminal half region of PCMD-1 (aa 1–242), which contains predicted coiled-coil and disordered regions (Fig. 3 E).

Co-localization of PCMD-1 and the SPD-5(CL) domain was confirmed by immunostaining using anti PCMD-1 and anti-GFP. In the absence of endogenous SPD-5, both GFP::SPD-5(CL) and GFP::SPD-5(CL+PA) co-localized with PCMD-1 on the centriole, whereas GFP::SPD-5(PA) did not (Fig. 3 F). From these results, we concluded that the direct interaction of the CL domain with PCMD-1 was required for centriolar localization of SPD-5.

Additionally, a yeast two-hybrid screen using PCMD-1 as the bait identified centrosomal proteins TAC-1, PLK-1 and PLK-2 (Fig. S2).

## Discussion

### A model for how SPD-5 proteins assemble into the mitotic PCM scaffold

We propose a model for SPD-5 scaffold formation during centrosome maturation (Fig. 4). SPD-5 is recruited to the centrosomes through the interaction between its CL domain and PCMD-1, which is localized to the centrioles (Erpf et al., 2019). (ii) As PCMD-1 interacts with both SPD-5 and PLK-1, this interaction promotes phosphorylation of the PReM domain of SPD-5 by PLK-1 in the vicinity of the centrioles. The phosphorylated PReM domain interacts with the PA domain of its own and/or other SPD-5 molecules. (iii) This interaction between phosphorylated PReM domains and the PA domains promotes SPD-5 multimerization and release from the vicinity of centrioles to expand the PCM scaffold.

**Figure 4 :**
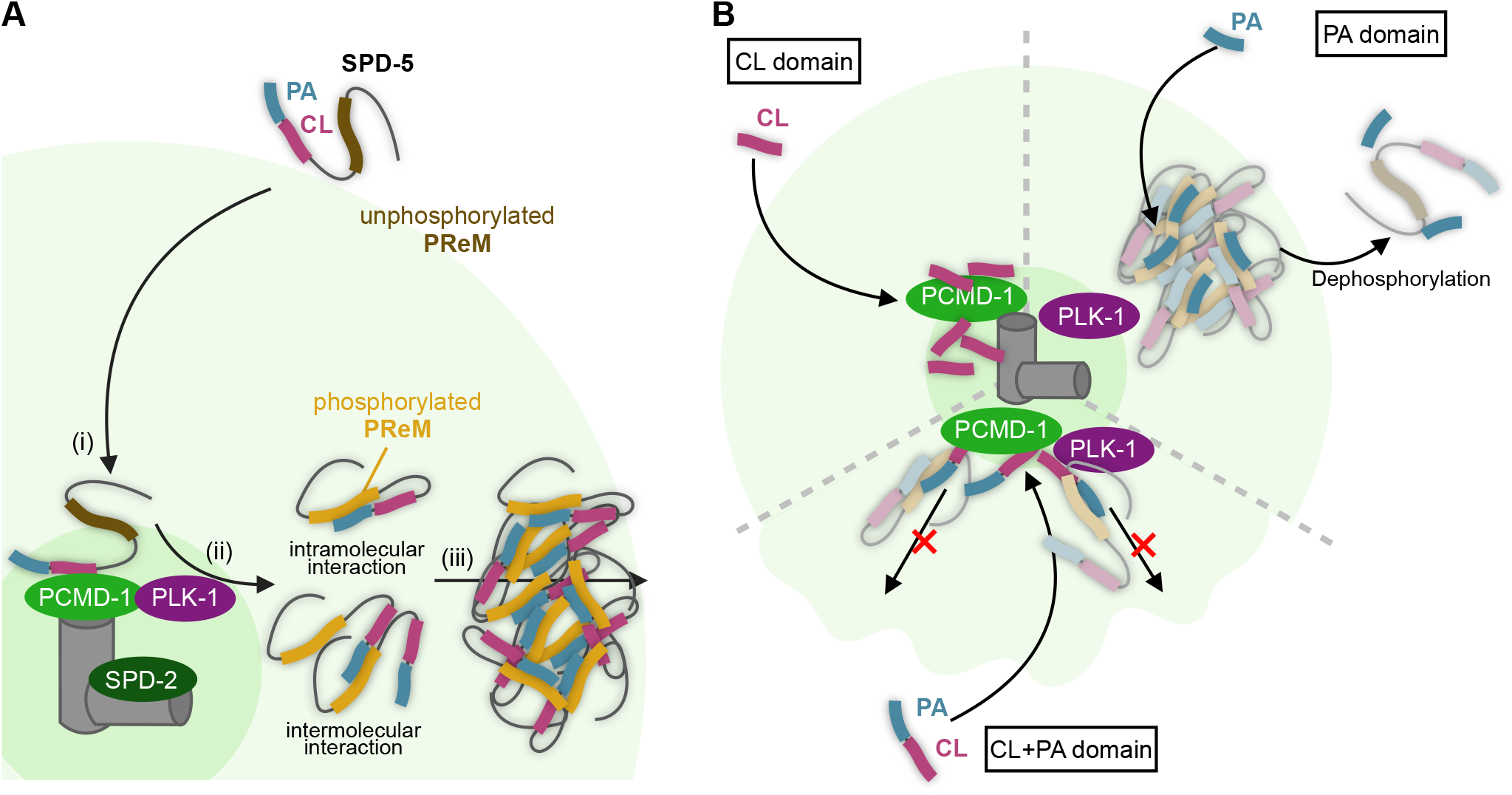
A model for PCM scaffold assembly by SPD-5. (A) model of PCM scaffold assembly by SPD-5 through PLK-1 phosphorylation dependent interactions. See text for details about steps i–iii. (B) Interpretation of the phenotypes of the CL, PA and CL+PA fragments observed in our *in vivo* analysis. The CL fragment localizes to centrioles through the interaction with centriolar PCMD-1. The PA fragment interacts with the phosphorylated PReM domain of endogenous SPD-5 and localizes to the central region of PCM. The CL+PA fragment ectopically anchors the phosphorylated endogenous SPD-5 to the proximity of centrioles, which prevents scaffold expansion and causes the dominant-negative effect.

Biochemically, the PA domain and the phosphorylated PReM domain interacted directly, and GFP::SPD-5(PA) localized to the PCM through intermolecular interactions with the PReM domain of endogenous SPD-5. However, PA-lacking GFP::SPD-5(ΔC3) localized to the centrioles but did not expand to the PCM, even though it contains the PReM domain, raising the possibility that the intramolecular interaction between the phosphorylated PReM and the PA domain is necessary for the incorporation of SPD-5 into the PCM. Based on these results, we speculate that both intramolecular and intermolecular interactions between SPD-5 regions are necessary for PCM formation (Fig. 4 A (ii)).

The CL+PA fragment (C1) caused a dominant-negative effect on the PCM expansion by endogenous SPD-5. Because this fragment interacts with both PCMD-1 and the PReM domain, it is likely to anchor endogenous SPD-5 ectopically within proximity of the centrioles, which prevents scaffold expansion (Fig. 4 B). In contrast, GFP::SPD-5(C-half) did not show this dominant-negative effect, even though it also contains the CL and PA domains. This difference suggests that at least part of the PReM domain contained in the C-half region is required for dissociation of the CL domain from PCMD-1 and further implicates intramolecular interactions between the PReM and PA domains in the process of PCM expansion.

Previous *in vitro* experiments suggested that SPD-5 plays a crucial role in PCM scaffold formation in *C. elegans*, which is dependent on phosphorylation-regulated interactions among coiled-coil proteins (Woodruff et al., 2017; Woodruff et al., 2015), but involvement of specific domains has not been reported. Our results indicated that PLK-1 phosphorylation promotes specific interactions between the PReM and the PA domains of SPD-5 that are crucial for scaffold formation. Thus, PCM scaffold formation is not solely dependent on non-specific interactions between coiled-coil proteins, but rather it requires well-organized, specific interactions.

### Evolutionarily conserved mechanisms of PCM scaffold formation

SPD-5 has been thought to be a functional homolog of *Drosophila* Centrosomin (Cnn) and human CDK5RAP2, although sequence homology is undetectable (Fong et al., 2008; Hamill et al., 2002; Megraw et al., 1999).

Despite the lack of sequence similarity, Cnn and SPD-5 have some common features in their domain structures. Their N termini have a binding domain for a γ-tubulin complex (Ohta et al., 2021; Zhang and Megraw, 2007). Both proteins have a domain for centrosomal targeting at their C terminus (CM2 for Cnn and CL for SPD-5) (Feng et al., 2017) and have a PReM domain in their central region that contains Polo/PLK-1 phosphorylation sites (Conduit et al., 2014a; Woodruff et al., 2015). Phosphorylation in the PReM domain by Polo/PLK-1 triggers the interaction between the PReM domain and a C-terminally located domain (CM2 for Cnn and PA for SPD-5), which leads to PCM scaffold assembly (Feng et al., 2017). Notably, the CM2 domain in Cnn carries out the combined roles of the CL and PA domains of SPD-5.

In addition to the similarity of the domain organization in Cnn and SPD-5, our results suggest that the spatial organization of centrosomes during scaffold assembly is also conserved between *Drosophila* and *C. elegans*. Cnn proteins are first recruited to the centrioles, where they are phosphorylated by Polo (Conduit et al., 2010; Conduit et al., 2014a). The phosphorylated Cnn proteins are believed to move away from the centrioles and eventually become dephosphorylated at the periphery of the PCM (Conduit et al., 2014a; Feng et al., 2017). Based on the more restricted centrosomal signal area of the PA fragment (GFP::SPD-5(C2)) in the presence of endogenous SPD-5, we speculate that phosphorylated SPD-5 proteins accumulate in the central part of the centrosomes. It has been shown that SPD-5 scaffold disassembly is mediated by dephosphorylation of SPD-5 (Enos et al., 2018; Mittasch et al., 2020). The dominant-negative effect of the CL+PA fragment of SPD-5 is also consistent with our model, as it proposes that the release of SPD-5 from the centrioles is required for proper PCM expansion. Thus, the spatial organization of the PCM seems to be conserved between *Drosophila* and *C. elegans*, in which the scaffold proteins are phosphorylated in the vicinity of centrioles, released, and eventually dephosphorylated in the periphery of the PCM.

In conclusion, we have revealed the PCM scaffold assembly process mediated by SPD-5 through *in vivo* domain analyses: SPD-5 is recruited to the centrioles through direct interaction between its CL domain and PCMD-1. Phosphorylation by centriolar PLK-1 of the PReM domain of SPD-5 promotes its intramolecular and/or intermolecular interactions with the PA domain, which leads to mitotic PCM scaffold assembly. Our findings further suggest an evolutionarily conserved domain organization between Cnn and SPD-5, and their phosphorylation-dependent regulation. Further analysis of SPD-5 and other PCM proteins will lead to a better understanding of the universality and diversity of the PCM formation mechanisms.

## Acknowledgments

We thank Yuji Kohara (National Institutes of Genetics, Mishima, Japan) for providing cDNA clones and the members of the Sugimoto laboratory for discussions. This work was supported by JSPS KAKENHI Grant Numbers JP15H04369 and JP15K14503 and a Bilateral Joint Research Project for A.S.; JP16K07334 and 20K06616 for N.H.; MEXT/JSPS WISE Program: Advanced Graduate Program for Future Medicine and Health Care, Tohoku University for M.N. The authors declare no competing financial interests.

## Author Contributions

Conceptualization, Hikaru Kano, Nami Haruta and Asako Sugimoto; Methodology, Momoe Nakajo, Hikaru Kano, Kenji Tsuyama, Nami Haruta and Asako Sugimoto; Investigation, Momoe Nakajo and Hikaru Kano; Writing – Original Draft, Momoe Nakajo; Writing – Review & Editing, Nami Haruta and Asako Sugimoto; Funding Acquisition, Momoe Nakajo, Nami Haruta and Asako Sugimoto

## Materials and methods

### Worm strains and culture

All *C. elegans* strains were cultured as described (Brenner, 1974). The strains used in this article are listed in the Key Resources Table and were maintained at 23°C. Worm strains used in this study are shown in Table1.

**Table 1:**
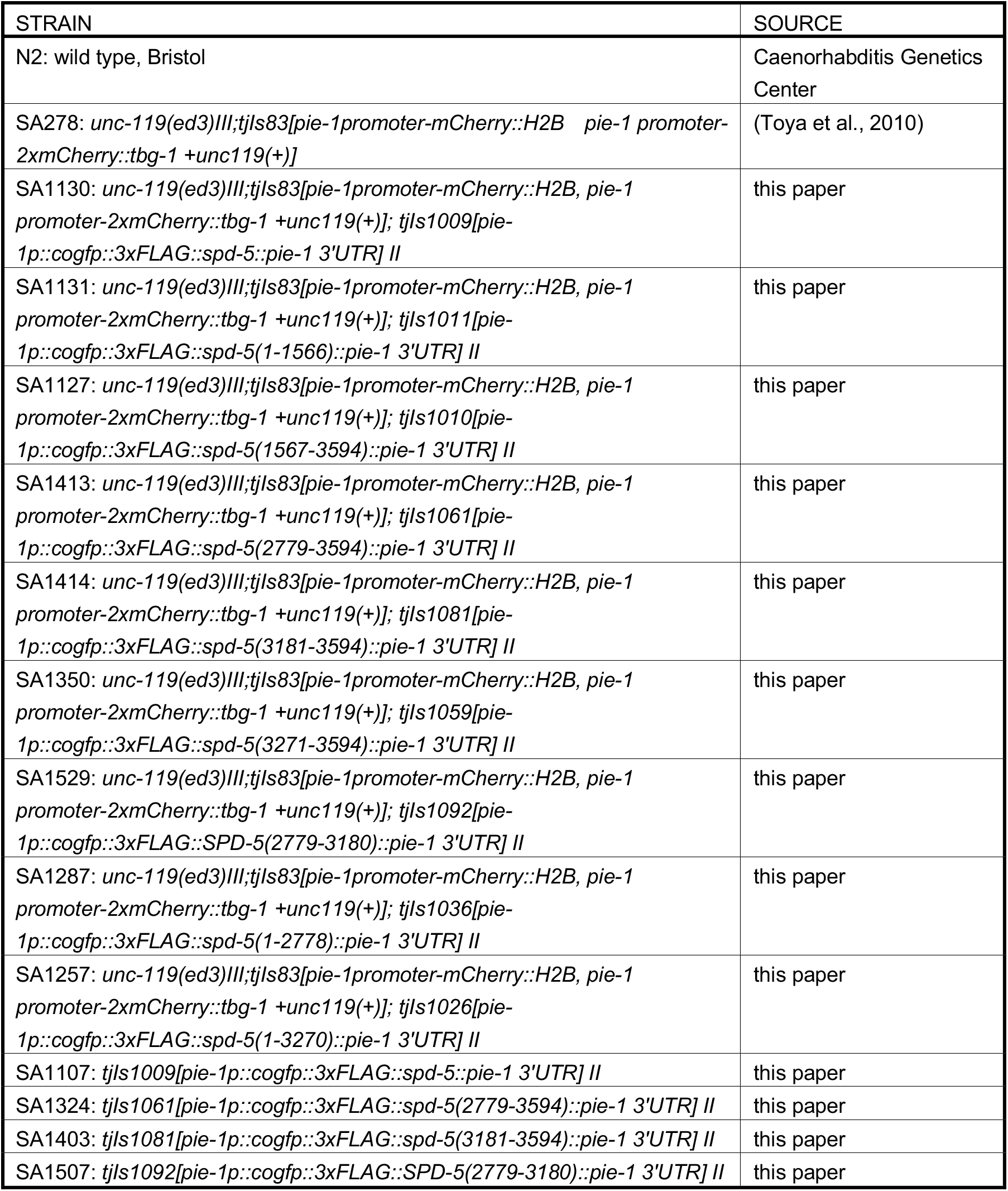
*C. elegans* strain list

| STRAIN | SOURCE |
| --- | --- |
| N2: wild type, Bristol | Caenorhabditis Genetics Center |
| SA278: *unc-119(ed3)III;tjls83[pie-1promoter-mCherry::H2B pie-1 promoter-2xmCherry::tbg-1 +unc119(+)]* | (Toya et al., 2010) |
| SA1130: *unc-119(ed3)III;tjls83[pie-1promoter-mCherry::H2B, pie-1 promoter-2xmCherry::tbg-1 +unc119(+)]*; *tjls1009[pie-1p::cogfp::3xFLAG::spd-5::pie-1 3'UTR] II* | this paper |
| SA1131: *unc-119(ed3)III;tjls83[pie-1promoter-mCherry::H2B, pie-1 promoter-2xmCherry::tbg-1 +unc119(+)]*; *tjls1011[pie-1p::cogfp::3xFLAG::spd-5(1-1566)::pie-1 3'UTR] II* | this paper |
| SA1127: *unc-119(ed3)III;tjls83[pie-1promoter-mCherry::H2B, pie-1 promoter-2xmCherry::tbg-1 +unc119(+)]*; *tjls1010[pie-1p::cogfp::3xFLAG::spd-5(1567-3594)::pie-1 3'UTR] II* | this paper |
| SA1413: *unc-119(ed3)III;tjls83[pie-1promoter-mCherry::H2B, pie-1 promoter-2xmCherry::tbg-1 +unc119(+)]*; *tjls1061[pie-1p::cogfp::3xFLAG::spd-5(2779-3594)::pie-1 3'UTR] II* | this paper |
| SA1414: *unc-119(ed3)III;tjls83[pie-1promoter-mCherry::H2B, pie-1 promoter-2xmCherry::tbg-1 +unc119(+)]*; *tjls1081[pie-1p::cogfp::3xFLAG::spd-5(3181-3594)::pie-1 3'UTR] II* | this paper |
| SA1350: *unc-119(ed3)III;tjls83[pie-1promoter-mCherry::H2B, pie-1 promoter-2xmCherry::tbg-1 +unc119(+)]*; *tjls1059[pie-1p::cogfp::3xFLAG::spd-5(3271-3594)::pie-1 3'UTR] II* | this paper |
| SA1529: *unc-119(ed3)III;tjls83[pie-1promoter-mCherry::H2B, pie-1 promoter-2xmCherry::tbg-1 +unc119(+)]*; *tjls1092[pie-1p::cogfp::3xFLAG::SPD-5(2779-3180)::pie-1 3'UTR] II* | this paper |
| SA1287: *unc-119(ed3)III;tjls83[pie-1promoter-mCherry::H2B, pie-1 promoter-2xmCherry::tbg-1 +unc119(+)]*; *tjls1036[pie-1p::cogfp::3xFLAG::spd-5(1-2778)::pie-1 3'UTR] II* | this paper |
| SA1257: *unc-119(ed3)III;tjls83[pie-1promoter-mCherry::H2B, pie-1 promoter-2xmCherry::tbg-1 +unc119(+)]*; *tjls1026[pie-1p::cogfp::3xFLAG::spd-5(1-3270)::pie-1 3'UTR] II* | this paper |
| SA1107: *tjls1009[pie-1p::cogfp::3xFLAG::spd-5::pie-1 3'UTR] II* | this paper |
| SA1324: *tjls1061[pie-1p::cogfp::3xFLAG::spd-5(2779-3594)::pie-1 3'UTR] II* | this paper |
| SA1403: *tjls1081[pie-1p::cogfp::3xFLAG::spd-5(3181-3594)::pie-1 3'UTR] II* | this paper |
| SA1507: *tjls1092[pie-1p::cogfp::3xFLAG::SPD-5(2779-3180)::pie-1 3'UTR] II* | this paper |

### Worm strain construction

Transgenes for GFP-tagged SPD-5 variants were inserted into the same genomic position on chromosome II (MosSCI *ttTi5605*) (Frøkjær-Jensen et al., 2012) by CRISPR/Cas9-mediated genome editing. We constructed a homologous repair template vector, pHK6, to insert SPD-5 variant transgenes into *ttTi5605*. pHK6 contains arms for homologous recombination at *ttTi5605* (Chr. II armL, 920 bp and armR, 967 bp); a GFP-encoding sequence under the control of the germline-specific *pie-1* promoter and 3’ untranslated region (3’UTR); the self-excising cassette (SEC) (Dickinson et al., 2015) for hygromycin B selection and the *ccdB* sequence, which was substituted by each SPD-5 variant sequence (pHK6: Chr. II armL_*pie-1*p_coGFP_SEC_3xFLAG_*ccdB*_*pie-1* 3′UTR_Chr. II armR). The SEC_3xFLAG_*ccdB* fragment was from pDD282 (Addgene #66823), except that the *let-858* 3’UTR was substituted by the *par-5* 3’UTR to prevent gene silencing in the germline. To construct homologous repair templates for SPD-5 variants, pHK6 was digested with ApaI restriction enzyme, which recognizes sites at both ends of the *ccdB* sequence, and the coding sequence for individual SPD-5 variants was assembled using the Gibson assembly method (Gibson et al., 2009).

For single guide RNA (sgRNA) expression, we constructed the pTK73 vector, which was modified from pRB1017 (Fire et al., 1998) to incorporate the sgRNA^(F+E)^ sequence (Chen et al., 2013). Then, the target sequence for the sgRNA (5’-GATATCAGTCTGTTTCGTAA-3’) at *ttTi5605* was inserted into pTK73 (pTK73_ ttTi5605).

For the transgene insertion by CRISPR/Cas9, the following injection mixture was injected into N2 worms: the homologous repair templates (pHK6_spd-5 fragments, 10 ng/μL); the sgRNA plasmid (pTK73_ ttTi5605, 50 ng/μL); the Cas9-expression plasmid pDD162 (Peft-3::Cas9, 50 ng/μL, Addgene #47549) (Dickinson et al., 2013) and the three injection markers pCFJ90 (Pmyo-2::mCherry, Addgene #19327), pCFJ104 (Pmyo-3::mCherry, Addgene #19328) and pGH8 (Prab-3::mCherry, Addgene #19359) (Frøkjaer-Jensen et al., 2008). Hygromycin B-resistant Roller worms that lacked mCherry fluorescence were selected as integrant candidates, and then were heat-shocked to remove the SEC. The integration and selection cassette excision were confirmed by sequencing.

Strain SA278 (mCherry::TBG-1; mCherry::Histone H2B) was constructed by microparticle bombardment(Toya et al., 2010). Dual-color strains (GFP::SPD-5(xx); mCherry::TBG-1; mCherry::Histone H2B) were constructed by crossing.

### RNA interference (RNAi)

For the depletion of endogenous SPD-5 by RNAi, we used the sequence corresponding to the *spd-5* 3’UTR or an N-terminal open reading frame (ORF) region (1–1566 bp). Template DNA fragments for synthesizing dsRNAs were amplified by PCR. The dsRNA was synthesized *in vitro* with T7 RiboMAX Express Large Scale RNA Production System (Promega, Madison, USA) and then was purified by phenol-chloroform extraction. RNAi was carried out using the soaking method(Maeda et al., 2001). Briefly, 10 early fourth larval stage (L4) worms were soaked in 2 mg/ml dsRNA solution and were incubated at 24.5°C for 24 h. Adult worms were then recovered from the dsRNA solution and incubated at 23°C on nematode growth medium (NGM) plates for 14–24 h before the worms were used for live imaging.

### Microscopy

Fluorescence images were obtained as described(Honda et al., 2017), with an Orca-R2 digital CCD camera (Hamamatsu Photonics, Shizuoka, Japan) mounted on an IX71 microscope (Olympus, Tokyo, Japan) using a CSU-X1 spinning disc confocal system (Yokogawa Electric Corporation, Tokyo, Japan). MetaMorph software (Molecular Devices, California, USA) was used to control the microscopes. Live images of one-cell embryos were obtained using a UPlanSApo 60×/1.30 silicone oil objective lens (Olympus) with camera gain 255 and without binning, with 40-second intervals, 15 *z*-slices at intervals of 1 μm and a 300-ms exposure for GFP and 1000-ms exposure for mCherry. Immunofluorescence images were obtained using a UPlanSApo 100×/1.40 oil objective lens (Olympus) with camera gain 0 and without binning; with 20 *z*-slices at intervals of 0.5 μm and 250-ms exposure for Alexa488, 100-ms exposure for Alexa568 and 1000-ms exposure for DAPI. ImageJ/Fiji software (NIH; https://imagej.net/Fiji) was used for processing and analyzing the obtained images.

### Immunostaining

To generate antibodies against SPD-5 and PCMD-1, full-length recombinant SPD-5 or a PCMD-1 peptide (CDEGFDSSSLKNNPASLQRD; aa 6–24) was injected into rabbits as per standard protocols (MBL Life Science and Cosmo Bio, respectively).

For immunostaining, embryos were freeze-cracked and fixed with −20°C methanol for 5 min and washed in PBS-T (0.05% Tween 20 in phosphate-buffered saline). After being incubated for 30 min in PBS-T containing 0.1% bovine serum albumin (BSA) for blocking, the slides were incubated with rat anti-GFP (1:200, diluted with PBS-T containing 0.1% BSA; Nacalai Tesque, cat#04404-84) and rabbit anti–SPD-5 (1:10,000; this study) or rabbit anti–PCMD-1 (1:10,000; this study) at 4°C overnight. Slides were washed with PBS-T and then were incubated with Alexa Fluor 488–conjugated donkey anti–rat IgG (1:500; Invitrogen, California, USA; RRID AB_2535794), Alexa Flour 594–conjugated donkey anti–rabbit IgG (1:500; Invitrogen; RRID AB_141637) and DAPI (1 mg/ml, 1:500; Dojinbo). After a 2-h incubation at room temperature, the slides were washed with PBS-T and mounted with ProLong Diamond (Thermo Fisher Scientific).

### Scoring embryonic lethality

To score embryonic lethality, L4 worms were transferred to NGM plates with OP50 and incubated at 23°C for 24 h. Each 1-day-old adult was then transferred to a new plate and was allowed to lay eggs for 6 h before being removed. The plates with eggs were incubated at 23°C for 24 h, and the number of eggs and larvae was counted. The embryonic lethality was calculated as the number of eggs out of the total number of eggs and larvae.

### PLK-1 purification

PLK-1 expression in Sf9 insect cells and its purification were performed as described (Woodruff and Hyman, 2015). PLK-1::6xHis was expressed in Sf9 cells, and the cells were harvested 72 h after the infection of baculovirus geverated by the Bac-to-Bac system (Invitrogen, cat#10359016). For purification, a HisTALON Superflow Cartridge (Takara Bio, Shiga, Japan) and AKTA start system (GE Healthcare, Illinois, USA) were used. PLK-1::6xHis was eluted with 80 mM imidazole, and the buffer was then exchanged to imidazole-free buffer using PD-10 Columns (GE Healthcare). Proteins were concentrated using 30K Amicon Ultra Centrifugal Filters (Merck, Darmstadt, Germany), flash-frozen in liquid nitrogen and stored at −80°C.

### Pull-down assay

To express the GST-tagged SPD-5(272–732_WT) or SPD-5(272–732_4A) in *E. coli*, a cold-induced promoter (pCold, Takara Bio) was used. The subcloned plasmids were transformed into *E. coli* BL21-CodonPlus (DE3) (Agilent Technologies, California, USA). 2mL of transformed *E. coli* preculture were added to 25 mL of LB medium containing 50 μg/ml Ampicillin and cultured for 3h at 37°C, followed by addition of 50 μL of 1M IPTG and culture for 8h at 16°C. Flash-frozen 50-μL bacterial pellets were dissolved in 500 μL binding buffer (PBS with 0.5 mM EDTA, 1 mM DTT, 1 mM PMSF, 26.3 μM MG-132 [Chemscene, New Jersey, USA], 2 cOmplete EDTA-free protease inhibitors [Roche, Basel, Switzerland] and 1× PhosSTOP [Roche]) and sonicated (1 min, six times) before centrifugation (21,000 ×g, 20 min, 4°C). The supernatant was filtered using a Millex-HV filter (0.45 μm; Merck) and incubated with 50 μL Glutathione Sepharose 4B beads (GE Healthcare) for 2 h at 4°C. Beads were washed three times with wash buffer (PBS with 1.2 M NaCl, 0.5 mM EDTA, 1 mM DTT, 1 mM PMSF, 26.3 μM MG-132, 2× cOmplete EDTA-free protease inhibitors and 1× PhosSTOP).

To prepare PLK-1 phosphorylated SPD-5 fragments, the Glutathione Sepharose 4B beads bound to GST-SPD-5(272–732) were incubated with 12 nM purified PLK-1 in reaction buffer (25 mM HEPES, pH 7.4; 150 mM KCl; 0.5 mM DTT; 0.2 mM ATP; 10 mM MgCl_2_; 0.025 mg/mL ovalbumin) (modified from (Woodruff and Hyman, 2015)). After a 40-min incubation at room temperature, the beads were washed three times with worm lysis buffer (PBS with 0.4 M NaCl, 0.5 mM EDTA, 1 mM DTT, 1 mM PMSF, 26.3 μM MG-132, 2× cOmplete EDTA-free protease inhibitors).

The beads bound to GST-SPD-5(272–732) with or without phosphorylation were added to the egg extract and incubated at 4°C for 2 h. For the preparation of egg extracts, ∼50 μL of flash-frozen eggs was added to 400 μL worm lysis buffer and sonicated (20 sec, six times) before centrifugation (21,000 × g, 15 min, 4°C). The beads were washed three times with worm lysis buffer, SDS-PAGE sample buffer (62.5 mM Tris-HCl, pH 6.8; 10% glycerol; 5% 2-mercaptoethanol; 2.5% SDS) was added and the sample was boiled at 98°C for 5 min. The samples were analyzed by western blotting.

### Western blotting

To check the expression level of transgenes, 15 one-day adult worms were collected in 10 μL of distilled water and flash-frozen in liquid nitrogen. The sample was then thawed and mixed with 3 μL of 4× SDS-PAGE sample buffer and 1.2 μL of 10% NP-40 and was boiled at 98°C for 5 min.

The samples were loaded onto a 5–20% gradient SDS-PAGE gel (Wako, Osaka, Japan) and transferred to an Immobilon-P Transfer Membrane (Merck). Antibodies were diluted with Signal Enhancer HIKARI for Western Blotting and ELISA (Nacalai Tesque). Primary antibodies rabbit anti-SPD-5 (1:2000) and rat anti-GFP (1:10,000) were generated against full-length recombinant proteins (MBL Life Science). Additional primary antibodies were mouse anti-GST B-14 (1:2000; Santa Cruz Biotechnology, Texas, USA; cat#sc-138) and mouse anti-α-tubulin DM1A (1:2000; Sigma-Aldrich, Missouri, USA; cat#T9026). Secondary antibodies were horseradish peroxidase (HRP)-conjugated goat anti-mouse IgG (H+L) (1:50,000; Jackson ImmunoResearch Laboratories, Pennsylvania, USA; RRID AB_2338511), HRP-conjugated goat anti-rabbit IgG (H+L) (1:50,000; Jackson ImmunoResearch Laboratories; RRID AB_10015282) and HRP-conjugated goat anti-rat IgG (H+L) (1:50,000; Jackson ImmunoResearch Laboratories; RRID AB_2340639). Signals were detected using Chemi-Lumi One Ultra (Nacalai Tesque), and blot images were obtained with a ChemiDoc Touch MP (Bio-Rad, California, USA).

### Yeast two-hybrid assay and screen

The ProQuest Two-Hybrid System (Invitrogen) was used for the yeast two-hybrid assay. Protein coding sequences were integrated into the plasmids pDEST22 and pDEST32, downstream of the GAL4 DNA activation domain and DNA binding domain, respectively. Yeast strain Mav203 (Invitrogen, cat#PQ1000101) was treated with Frozen-EZ Yeast Transformation II (Zymo Research, California, USA) to generate competent cells and was then transformed with bait and prey plasmids. The transformants were selected by culturing on SC-WL plates for 3 days at 30°C. Each yeast strain was cultured in 2 mL of liquid SC-WL for 13 h at 30°C and was analyzed on selection plates containing 3-amino-1,2,4-triazole (3AT).

For the yeast two-hybrid screen, we first transformed yeast Mav203 with bait plasmids (the coding sequence of SPD-5 fragments or PCMD-1 subcloned into pDEST32) only and incubated the cells on SC-W plates. The competent yeast cells with a single bait vector were then used in a transformation with the prey vector (cDNA fragments cloned into a pPC86 plasmid, Invitrogen) for yeast two-hybrid screening. Transformation efficiency was measured based on the number of colonies on SC-WL plates. For the screening, transformants were spread onto 20 SC-WL + 25 mM 3AT selection plates and incubated for 5–7 days at 30°C. Individual colonies were then picked and spread on new SC-WL plates. As candidate interactors with SPD-5(CL), 21 colonies were obtained from 1.9 × 10^5^ cells on selection plates. As candidate interactors with PCMD-1, 12 colonies were obtained from 6.6 × 10^4^ cells on selection plates. The coding sequences of the candidate clones were PCR amplified and sequenced (Forward primer: TATAACGCGTTTGGAATCACT, Reverse primer: AGCCGACAACCTTGATTGGAGAC), and their identities were determined by a WormBase BLAST/BLAT search (https://wormbase.org/tools/blast_blat). For all positive prey plasmids, interactions were confirmed by isolation from the positive colonies and re-transformation to Mav 203 with the bait vector.

## Statistical analysis

The details for quantification, sample numbers and statistical analyses are described in the main text, the figures and the figure legends. The values given for n represent the number of embryos imaged to determine phenotypes (Fig. 1 A–C) or the number of embryos counted to measure embryonic lethality (Fig. 2 B).

GraphPad Prism 7 software was used to analyze and represent the data. Results are expressed as the mean value ± SEM (Fig. 1 B) or the mean value + SEM (Fig. 2 B). A one-way ANOVA and Tukey’s multiple comparisons test were performed to determine significant differences (Fig. 1 B and 2 B). Significance levels: n.s., p ≥ 0.05, *p < 0.05, **p < 0.01, ***p < 0.001.

The centrosomal GFP signal was labeled using the threshold set by the Yen algorithm (Fiji/ImageJ), and its area was measured (Fig. 1 B).

## References

Alvarez-Rodrigo, I., T.L. Steinacker, S. Saurya, P.T. Conduit, J. Baumbach, Z.A. Novak, M.G. Aydogan, A. Wainman, and J.W. Raff. 2019. Evidence that a positive feedback loop drives centrosome maturation in fly embryos. Elife. 8.

Barr, A.R., J.V. Kilmartin, and F. Gergely. 2010. CDK5RAP2 functions in centrosome to spindle pole attachment and DNA damage response. J Cell Biol. 189:23–39.

Brenner, S. 1974. The genetics of Caenorhabditis elegans. Genetics. 77:71–94.

Chen, B., L.A. Gilbert, B.A. Cimini, J. Schnitzbauer, W. Zhang, G.W. Li, J. Park, E.H. Blackburn, J.S. Weissman, L.S. Qi, and B. Huang. 2013. Dynamic imaging of genomic loci in living human cells by an optimized CRISPR/Cas system. Cell. 155:1479–1491.

Conduit, P.T., K. Brunk, J. Dobbelaere, C.I. Dix, E.P. Lucas, and J.W. Raff. 2010. Centrioles regulate centrosome size by controlling the rate of Cnn incorporation into the PCM. Curr Biol. 20:2178–2186.

Conduit, P.T., Z. Feng, J.H. Richens, J. Baumbach, A. Wainman, S.D. Bakshi, J. Dobbelaere, S. Johnson, S.M. Lea, and J.W. Raff. 2014a. The centrosome-specific phosphorylation of Cnn by Polo/Plk1 drives Cnn scaffold assembly and centrosome maturation. Dev Cell. 28:659–669.

Conduit, P.T., J.H. Richens, A. Wainman, J. Holder, C.C. Vicente, M.B. Pratt, C.I. Dix, Z.A. Novak, I.M. Dobbie, L. Schermelleh, and J.W. Raff. 2014b. A molecular mechanism of mitotic centrosome assembly in Drosophila. Elife. 3:e03399.

Decker, M., S. Jaensch, A. Pozniakovsky, A. Zinke, K.F. O’Connell, W. Zachariae, E. Myers, and A.A. Hyman. 2011. Limiting amounts of centrosome material set centrosome size in C. elegans embryos. Curr Biol. 21:1259–1267.

Dickinson, D.J., A.M. Pani, J.K. Heppert, C.D. Higgins, and B. Goldstein. 2015. Streamlined Genome Engineering with a Self-Excising Drug Selection Cassette. Genetics. 200:1035–1049.

Dickinson, D.J., J.D. Ward, D.J. Reiner, and B. Goldstein. 2013. Engineering the Caenorhabditis elegans genome using Cas9-triggered homologous recombination. Nat Methods. 10:1028–1034.

Enos, S.J., M. Dressler, B.F. Gomes, A.A. Hyman, and J.B. Woodruff. 2018. Phosphatase PP2A and microtubule-mediated pulling forces disassemble centrosomes during mitotic exit. Biol Open. 7.

Erpf, A.C., L. Stenzel, N. Memar, M. Antoniolli, M. Osepashvili, R. Schnabel, B. Conradt, and T. Mikeladze-Dvali. 2019. PCMD-1 Organizes Centrosome Matrix Assembly in C. elegans. Curr Biol. 29:1324–1336.e1326.

Feng, Z., A. Caballe, A. Wainman, S. Johnson, A.F.M. Haensele, M.A. Cottee, P.T. Conduit, S.M. Lea, and J.W. Raff. 2017. Structural Basis for Mitotic Centrosome Assembly in Flies. Cell. 169:1078–1089.e1013.

Fire, A., S. Xu, M.K. Montgomery, S.A. Kostas, S.E. Driver, and C.C. Mello. 1998. Potent and specific genetic interference by double-stranded RNA in Caenorhabditis elegans. Nature. 391:806–811.

Fong, K.W., Y.K. Choi, J.B. Rattner, and R.Z. Qi. 2008. CDK5RAP2 is a pericentriolar protein that functions in centrosomal attachment of the gamma-tubulin ring complex. Mol Biol Cell. 19:115–125.

Frøkjaer-Jensen, C., M.W. Davis, C.E. Hopkins, B.J. Newman, J.M. Thummel, S.P. Olesen, M. Grunnet, and E.M. Jorgensen. 2008. Single-copy insertion of transgenes in Caenorhabditis elegans. Nat Genet. 40:1375–1383.

Frøkjær-Jensen, C., M.W. Davis, M. Ailion, and E.M. Jorgensen. 2012. Improved Mos1-mediated transgenesis in C. elegans. Nat Methods. 9:117–118.

Giansanti, M.G., E. Bucciarelli, S. Bonaccorsi, and M. Gatti. 2008. Drosophila SPD-2 is an essential centriole component required for PCM recruitment and astral-microtubule nucleation. Curr Biol. 18:303–309.

Gibson, D.G., L. Young, R.Y. Chuang, J.C. Venter, C.A. Hutchison, and H.O. Smith. 2009. Enzymatic assembly of DNA molecules up to several hundred kilobases. Nat Methods. 6:343–345.

Gomez-Ferreria, M.A., U. Rath, D.W. Buster, S.K. Chanda, J.S. Caldwell, D.R. Rines, and D.J. Sharp. 2007. Human Cep192 is required for mitotic centrosome and spindle assembly. Curr Biol. 17:1960–1966.

Hamill, D.R., A.F. Severson, J.C. Carter, and B. Bowerman. 2002. Centrosome maturation and mitotic spindle assembly in C. elegans require SPD-5, a protein with multiple coiled-coil domains. Dev Cell. 3:673–684.

Haren, L., T. Stearns, and J. Lüders. 2009. Plk1-dependent recruitment of gamma-tubulin complexes to mitotic centrosomes involves multiple PCM components. PLoS One. 4:e5976.

Honda, Y., K. Tsuchiya, E. Sumiyoshi, N. Haruta, and A. Sugimoto. 2017. Tubulin isotype substitution revealed that isotype combination modulates microtubule dynamics in. J Cell Sci. 130:1652–1661.

Kemp, C.A., K.R. Kopish, P. Zipperlen, J. Ahringer, and K.F. O’Connell. 2004. Centrosome maturation and duplication in C. elegans require the coiled-coil protein SPD-2. Dev Cell. 6:511–523.

Maeda, I., Y. Kohara, M. Yamamoto, and A. Sugimoto. 2001. Large-scale analysis of gene function in Caenorhabditis elegans by high-throughput RNAi. Curr Biol. 11:171–176.

Megraw, T.L., K. Li, L.R. Kao, and T.C. Kaufman. 1999. The centrosomin protein is required for centrosome assembly and function during cleavage in Drosophila. Development. 126:2829–2839.

Mittasch, M., V.M. Tran, M.U. Rios, A.W. Fritsch, S.J. Enos, B. Ferreira Gomes, A. Bond, M. Kreysing, and J.B. Woodruff. 2020. Regulated changes in material properties underlie centrosome disassembly during mitotic exit. J Cell Biol. 219.

Ohta, M., Z. Zhao, D. Wu, S. Wang, J.L. Harrison, J.S. Gómez-Cavazos, A. Desai, and K.F. Oegema. 2021. Polo-like kinase 1 independently controls microtubule-nucleating capacity and size of the centrosome. J Cell Biol. 220.

Palazzo, R.E., J.M. Vogel, B.J. Schnackenberg, D.R. Hull, and X. Wu. 2000. Centrosome maturation. Curr Top Dev Biol. 49:449–470.

Pelletier, L., N. Ozlü, E. Hannak, C. Cowan, B. Habermann, M. Ruer, T. Müller-Reichert, and A.A. Hyman. 2004. The Caenorhabditis elegans centrosomal protein SPD-2 is required for both pericentriolar material recruitment and centriole duplication. Curr Biol. 14:863–873.

Toya, M., Y. Iida, and A. Sugimoto. 2010. Imaging of mitotic spindle dynamics in Caenorhabditis elegans embryos. Methods Cell Biol. 97:359–372.

Woodruff, J.B., B. Ferreira Gomes, P.O. Widlund, J. Mahamid, A. Honigmann, and A.A. Hyman. 2017. The Centrosome Is a Selective Condensate that Nucleates Microtubules by Concentrating Tubulin. Cell. 169:1066–1077.e1010.

Woodruff, J.B., and A.A. Hyman. 2015. Method: In vitro analysis of pericentriolar material assembly. Methods Cell Biol. 129:369–382.

Woodruff, J.B., O. Wueseke, V. Viscardi, J. Mahamid, S.D. Ochoa, J. Bunkenborg, P.O. Widlund, A. Pozniakovsky, E. Zanin, S. Bahmanyar, A. Zinke, S.H. Hong, M. Decker, W. Baumeister, J.S. Andersen, K. Oegema, and A.A. Hyman. 2015. Centrosomes. Regulated assembly of a supramolecular centrosome scaffold in vitro. Science. 348:808–812.

Wueseke, O., D. Zwicker, A. Schwager, Y.L. Wong, K. Oegema, F. Jülicher, A.A. Hyman, and J.B. Woodruff. 2016. Polo-like kinase phosphorylation determines Caenorhabditis elegans centrosome size and density by biasing SPD-5 toward an assembly-competent conformation. Biol Open. 5:1431–1440.

Zhang, J., and T.L. Megraw. 2007. Proper recruitment of gamma-tubulin and DTACC/Msps to embryonic Drosophila centrosomes requires Centrosomin Motif 1. Mol Biol Cell. 18:4037–4049.

