## Supplementary material for "Centrosome maturation requires phosphorylation-mediated sequential domain interactions of SPD-5": Supplemenal Material

### Supplemental Information titles and legends

**Figure S1 : Expression levels of GFP-labeled truncated versions of SPD-5**

Western blots to check the expression levels of each GFP-labeled truncated version of SPD-5 (GFP::SPD-5(xx)). (See Fig. 1 A for a schematic of these truncated variants.) The red arrowheads indicate the band associated with each GFP-tagged truncated SPD-5 fragment. At the first lane, "–" indicate wild-type worms. Molecular weights were calculated as follows: GFP::SPD-5(FL), 166.9 kDa; GFP::SPD-5(N-half), 90.2 kDa; GFP::SPD-5(C-half), 108.5 kDa; GFP::SPD-5(C1), 62.9 kDa; GFP::SPD-5(C2), 44.3 kDa; GFP::SPD-5(C3), 47.6 kDa; GFP::SPD-5(C4), 47.2 kDa; GFP::SPD-5(ΔC1), 135.7 kDa; GFP::SPD-5(ΔC3), 154.5 kDa; endogenous SPD-5, 135.1 kDa; α-tubulin, ~50 kDa.

**Figure S2 : PCMD-1 interacts with centrosomal proteins TAC-1, PLK-1 and PLK-2**

The results of a yeast two-hybrid screen to identify PCMD-1–interacting proteins. The growth of the relevant strains on SC/−Leu/−Trp plates and SC/−Leu/−Trp/−Ura plates containing 100 mM 3-amino-1,2,4-triazol (3AT). C17E4.20 is a unnamed gene.

### Supplemental Figure 1

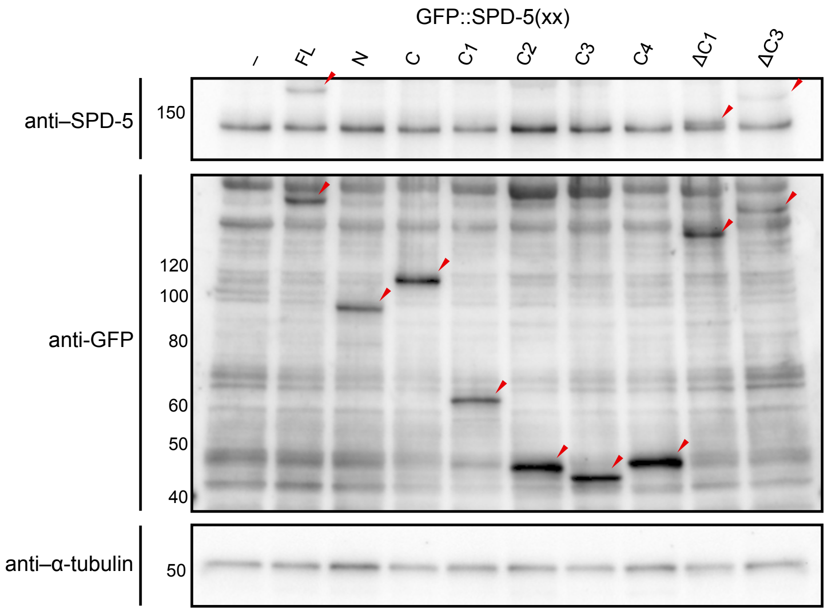

**Supplemental Figure 2**

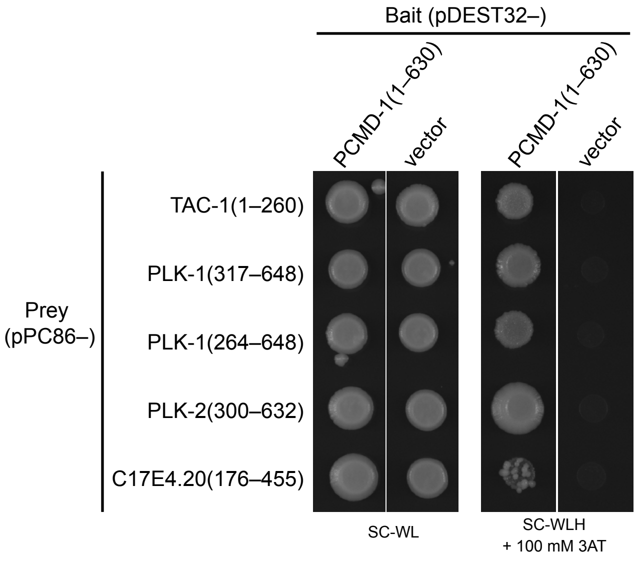
